# Scaling-up the production of recombinant EIT antigen by *E. coli* fermentation for a vaccine-preparation against EHEC for cattle

**DOI:** 10.64898/2025.12.13.694097

**Authors:** Laura Ana Basile, Diego Gabriel Noseda, Ingrid Milstein, Gabriel Briones

## Abstract

**Aims:** This study aims to optimize and scale up the production of the chimeric antigen EIT through *E. coli* fed-batch fermentations. The approach seeks for simplicity and cost-effectiveness, considering EIT as a potential vaccine candidate against EHEC for cattle. Its ability to induce a humoral immune response has been recently verified in bovines in a proof-of-concept study.

**Methods and Results:** An initial screening was conducted to select the optimal medium for EIT expression, using lactose for recombinant protein induction. M9 Minimal medium supplemented with yeast extract yielded the highest relative levels of EIT, as determined by Western blot analysis. *E. coli* cultures were subsequently grown in a stirred-tank bioreactor, and both biomass production and EIT expression were monitored. The process was reproducible across three independent fermentations, with all parameters being improved compared with shake-flask cultivations. Antigen recovery was achieved through thermal permeabilization, as the construction includes a periplasmic signal sequence. Overall, the process resulted in an average potential output of 350 doses per liter of fermented culture, while preserving the EIT antigenic capability, as confirmed by ELISA assays.

**Conclusion:** The production of the recombinant EIT antigen was successfully scaled up using a stirred-tank bioreactor through a quite simple and cost-effective approach, achieving increased yields for supporting further studies and interventions.

**Impact Statement:** Cattle are the major reservoir and source of dissemination of enterohemorrhagic *E. coli* (EHEC), a human pathogen responsible for outbreaks of bloody diarrhea and hemolytic uremic syndrome (HUS) worldwide. A scalable and cost-effective preharvest vaccine for cattle could help with developing strategies aimed at reducing bacterial carriage and thus, the impact of this zoonosis.

## Introduction

Enterohemorrhagic *Escherichia coli* (EHEC) is a human pathogen responsible for outbreaks of bloody diarrhea and hemolytic uremic syndrome (HUS) worldwide (Nguyen and Sperandio 2012). Being cattle a major natural reservoir and a recognized source of environmental dissemination, EHEC strains have been linked to numerous foodborne outbreaks (Bell 2002, Ferens and Hovde 2011, Greig and Ravel 2009). Strategies aimed at reducing EHEC carriage and shedding in cattle are key to reduce the impact of this zoonosis and thus, EHEC infections in humans.

EIT is a chimeric antigen previously designed as a vaccine candidate against EHEC (Amani et al. 2009). It is a trivalent recombinant protein composed of immunologically relevant regions of EspA, Intimin and Tir - proteins involved in the T3SS-dependent attachment of EHEC to the intestinal epithelium. EIT’s antigenic capability has been explored through different methodological approaches seeking for a vaccine development: production in plant systems for an oral EHEC vaccine (Amani et al. 2011); cytoplasmic production in *E. coli* BL21 followed by purification and incorporation into non-pathogenic bacteria or bacterial vaccine strains as delivery vehicles for oral administration (Iannino et al. 2015, 2022; Uriza et al. 2020); and linkage to chitosan nanoparticles for intranasal immunization (Doavi et al. 2016), among others. Across these approaches, EIT immunization has been shown to be effective in murine model, inducing robust humoral response and even reducing faecal shedding.

*E. coli* laboratory strain BL21 is one of the most widely used systems for recombinant protein production. Its main advantages include fast growth in simple and inexpensive media, high protein expression levels, the availability of strains with low proteolytic activity, and extensive scientific knowledge regarding its physiology and genetics (Ferreira et al. 2018).

In a recent work, we engineered EIT for expression and periplasmic localization in *E. coli* BL21 (Duarte et al. 2024). The antigen production was designed seeking for simplicity and cost-effectiveness, as these factors often pose challenges for the adoption of vaccination programs in the agricultural sector. In this approach, EIT synthesis is induced by lactose supplementation and a signal-peptide for periplasmic transport is incorporated to facilitate downstream processing. Following a thermal shock, the antigen-enriched periplasmic fraction was used to formulate a vaccine preparation for preharvest administration in cattle. As proof of concept, subcutaneous application elicited immunity in both mouse and bovine models, evidenced by ELISA assays and actin-pedestal inhibition in cell cultures (Duarte et al. 2024). In that initial assessment, the recombinant protein was produced in shake-flask *E. coli* cultures grown in LB media, at a scale large enough for process characterization and immunization assays but resulting in a limited number of doses. The aim of this work was to scale up *E. coli* cultivation to a stirred-tank bioreactor evaluating both biomass and antigen production.

Fermentation scale up aims to recover larger product quantities while maintaining -or ideally improving-specific yield and product quality, key issues in industrial production processes (Schmidt 2005). Searching for higher productivity with cost-effective media and short incubation times, we performed a first screening in replicated shake-flask cultures. By using a fed-batch strategy, three fermentation processes in a stirred-tank bioreactor were performed in the established conditions to analyse the reproducibility of the fermentation parameters and the optimal timing for downstream processing.

## Materials and methods

### 1. Strain and vector used for expression of the EIT recombinant antigen

*Escherichia coli* BL21 (DE3) strain was used for recombinant EIT expression. The recombinant EIT antigen has been already generated and characterized (Duarte et al. 2024). Briefly, the construction consists of the C-terminal region of EspA (EspA 36-192), the Tir-binding domain of Intimin (Int 653-935), and the Intimin-binding domain of Tir (Tir 258-361) (strain EDL933, i.e. serotype O157:H7), separated by a link sequence (EAAAK)_4_ to promote proper folding. The EIT chimeric sequence is under the control of the lactose/IPTG-inducible pTRC promoter. To direct EIT to the bacterial periplasm, the β-lactamase signal peptide (1-35) is fused to the N-terminus of the EIT open reading frame. The construction is carried on the pBBR1MCS-2 vector (Km^R^).

### 2. Optimization of culture media for EIT expression in shake flasks

The performance of EIT expression was comparatively evaluated by culturing *E. coli* transformant clone in shake flasks with LB (10 g triptone, 5 g yeast extract, and 5 g NaCl/L), M9 minimal media (1 g NH_4_Cl, 3 g KH_2_PO_4_, 6 g Na_2_HPO_4_, 0.5 g NaCl, 0.1 ml of 1M CaCl_2_, and 1 ml of 1 M MgSO_4_/L) and M9 supplemented with yeast extract at 0.1 % p/v (M9Y). Furthermore, the effect of addition of higher yeast extract content (0.3 % p/v) or trace metal elements (Studier 2005) to M9Y media was analysed. In all cases, media was supplemented with an auto-induction solution composed of 5 g glycerol, 0.5 g glucose, and 2 g lactose/L (referred as GGL5052) (Studier 2005). When necessary, kanamycin (Km) was added at a final concentration of 50 μg/ml.

Cultivations (20 ml) were initiated with an overnight EIT-carrying *E. coli* culture (dilution 1/50) performed in the corresponding medium without lactose. Cultures were grown at 37 °C and 200 rpm in an orbital shaker for 6 h. Periodically, OD at 600 nm was measured and 1 ml samples were collected for further analysis. The strain *E. coli* BL21 carrying the empty pBBR1MCS-2 vector was grown in LB as a control. Culture assays were performed at least in duplicate.

### 3. Fermentations in stirred-tank bioreactor

Fermentation of EIT-carrying *E. coli* clone was conducted in a 2.5 L stirred-tank bioreactor (EZ2-Control, Applikon, Getinge) applying a two-step process. The first phase consisted of a batch culture in M9Y medium supplemented with auto-induction solution (GGL5052), maintaining the temperature at 37 ± 0.1 °C and pH at 7.0 by automatically addition of 3 N HCl and 2.5 N NaOH. In this stage, the agitation was kept at 600 rpm and filtered (0.22 μm) compressed air was supplied at 1 VVM. Afterwards, a fed-batch stage was performed by feeding the culture with a solution containing 100 g/L glycerol, 40 g/L lactose, 20 g/L yeast extract and 3X M9 mineral salts. The feeding was carried out with a constant flow of 1 ml/min. In this phase, the dissolved oxygen was regulated near 30 % with stirring cascade (max: 750 rpm - min: 400 rpm) and by filtered compressed air (1 - 2 VVM). The stirred-tank bioreactor operated in interface with the *Lucullus* software (Applikon) for parameters control and data acquisition. The pH was measured with a pH electrode and dissolved oxygen level was determined by a polarographic probe (Applisens, Applikon). Foam formation during fermentation processes was avoided by addition of 4 % v/v antifoam agent 204 (Sigma-Aldrich). Fermentation was initiated by inoculating 1 L of M9Y-GGL5052 medium with 100 ml of an overnight *E. coli* culture grown in the same medium without lactose. Samples were withdrawn throughout the fermentation to monitor biomass (OD 600) and recombinant antigen expression. Two additional fermentations were performed under the same conditions to assess process reproducibility. In parallel with each fermentation, a 50 ml shake-flask culture was inoculated from the same starter culture and grown at 37 °C and 200 rpm to compare production parameters.

### 4. Expression analysis

Samples (1 ml) from both shake flask and bioreactor cultures were centrifuged at 3,500*xg* for 3 min, and biomass was resuspended in an appropriated volume of loading buffer. EIT expression was monitored by SDS-PAGE and Western blot analysis using the Odyssey Imaging System (LI-COR), with primary anti-EITH7 mouse antibodies and IR-Dye fluorophore-labeled secondary anti-mouse antibodies (LI-COR), as previously described (Uriza et al. 2020). ImageJ software (Integrated-Density measures) was used for relative or absolute -by using an EITH7 standard curve-quantitation of the detected bands. Biomass levels -converted to dry cell weight by using the approximation of 1 OD = 0.28 g/L (Hassan and Fridovich 1978)- and EIT quantities were used for production parameter calculations: i.e. EIT concentration (mg/L), yield of EIT relative to biomass formation (Y_p/x_, mg/g), volumetric productivity (P_v_, mg/L h) and specific productivity (P_s_, mg/g h).

### 5. Downstream processing: Thermal shock

After 7 h of each fermentation process, the culture was centrifuged at 8,000×*g* for 20 min at 4 °C in a Sorvall high-speed centrifuge (Lynx 4000 Thermo) for downstream processing. Pellet was incubated overnight at -20 °C. Thermal permeabilization of cells was carried out according to Tsuchido et al. (1985) with some modifications. Briefly, bacteria were suspended in TE buffer (50 mM Tris-HCl pH 7.6, 10 mM EDTA) by using around 4 % v/v of the original culture volume. The suspension was incubated on ice for 20 min, followed by a heat treatment at 55 °C for 30 min, using a water bath. Samples were immediately put in ice and centrifuged at 8,000× *g* for 25 min at 4 °C. Supernatants containing the periplasmic fraction were collected and stored at -20 °C for further analysis. EIT determination was carried out by Western blot as described above. For total protein profile, SDS-PAGE and Coomassie blue staining were performed. Samples were quantified by the Bradford method for total periplasmic protein content, by using a BSA standard curve.

### 6. Enzyme-Linked Immuno Sorbent Assay (ELISA)

The EIT recombinant antigen produced by *E. coli* fermentations was subjected to indirect ELISA. Briefly, samples of the periplasmic fraction in coating buffer were used to coat a 96 wells-microplate (150 ng EIT/well), using purified MBP-EITH7 in the control wells. Wells were incubated with pre- and post-immune sera from EIT-immunized bovines (Duarte et al. 2024) at a 1:300 dilution. Detection was performed using peroxidase-conjugated anti-bovine IgG (1:4000 dilution) and the peroxidase substrate TMB (tetramethylbenzidine, Sigma), according to established protocols. OD at 450 nm was determined in Tecan-Spark microplate reader. Samples were evaluated in triplicate; data were analysed by One-way ANOVA and Tukey contrasts in GraphPad software.

## Results

### 1. Selection of the optimal medium for EIT expression

Growth and antigen production were evaluated in LB, M9, and M9Y media in at least two independent experiments in shake flasks. In all cases, induction was achieved by auto-induction solution with lactose. This solution, referred as GGL5052, is a mixture of carbon sources in which glucose is used firstly for increasing biomass level while preventing basal expression, and then glycerol and lactose are intended to be used as the carbon source and inductor for recombinant protein expression, respectively (Studier 2005). EIT-producing *E. coli* cultures were incubated for 6 h; OD at 600 nm was measured and EIT expression was analysed for each condition by Western blot assays (Fig. 1). In addition to the full-length recombinant protein (65 kDa), smaller specific bands (around 50 and 37 kDa) were detected, likely resulting from proteolytic fragmentation or incomplete translation, as previously noted (Iannino et al. 2015).

**Figure 1.**
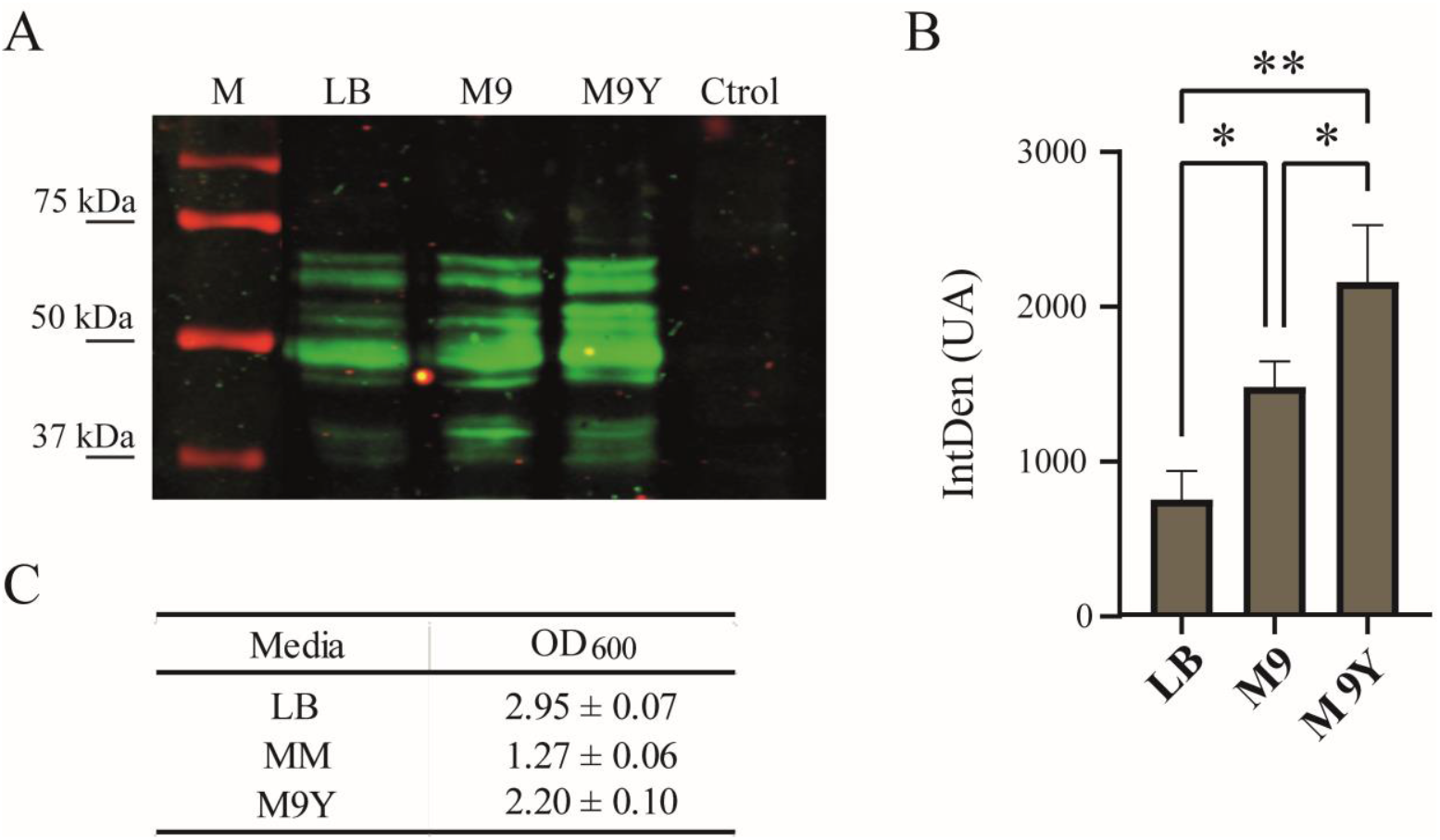
Selection of the optimal media for EIT expression. **A**. Representative Western blot image for EIT determination in LB, M9 and M9Y growth media. M: molecular weight marker, Ctrol: control (*E. coli* carrying the empty vector). **B**. Intensity of EIT bands in Western blot determined by ImageJ. Data correspond to two independent assays and were analysed by one-way ANOVA and Tukey contrasts. Asterisks indicate significant differences at p<0.05 (*) or p<0.01 (**). **C**. Growth in LB, M9, and M9Y media: OD600 values at 6 h of incubation are tabulated.

Despite the higher biomass level reached in LB, the relative level of EIT detected by Western blot duplicated in M9 minimal medium (M9) (Fig. 1). Yeast supplementation of M9 medium (M9Y) allowed an intermediate biomass level with the highest EIT expression (3- and 1.5-fold respect to LB and M9, respectively). Therefore, M9Y medium was selected for further fermentations. Addition of trace elements or higher yeast content (0.3 %) did not significantly enhance EIT production (data not shown).

### 2. Fermentation of *E. coli* BL21 expressing EIT antigen in a stirred-tank Bioreactor

Following selection of M9Y as the best medium for EIT expression, *E. coli* was grown in stirred-tank bioreactor using a two-step strategy: an initial 4-hour batch phase followed by a fed-batch step. Biomass production and EIT expression were monitored throughout the fermentation. Initially, the process was continued for 28 h (Fig. 2A and 2B left panel). The intensity profile of Western blot signals revealed that EIT levels increased during the first 7 h of fermentation, declining afterwards (Fig. 2A, bottom panel). The decrease in EIT in the late hours of fermentation could result from product degradation due to *E. coli* proteases or cell damage once exponential growth is no longer sustained, but this remains to be determined.

**Figure 2.**
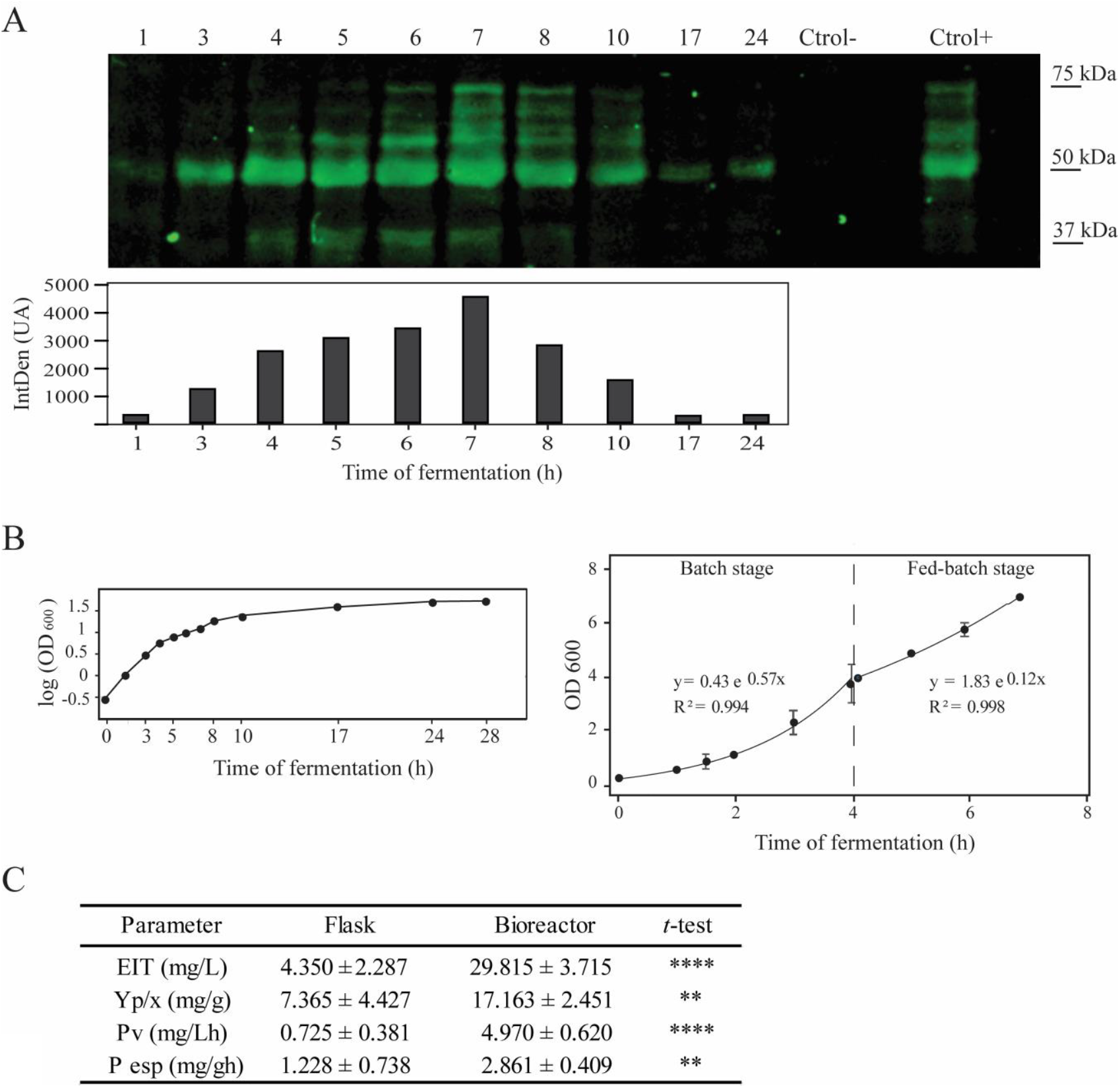
Fermentation of EIT-expressing *E. coli* BL21 in M9Y. **A**. EIT production along fermentation time (h) was evaluated by Western blot with specific anti-EIT primary antibodies. A representative image of one fermentation process is shown. Intensity profile of EIT bands as determined by ImageJ is plotted at the bottom panel. Samples were normalized to equal biomass prior to loading. Ctrol-refers to *E. coli* carrying the empty vector, Ctrol+ refers to a previous preparation of EIT. **B**. Bacterial growth in stirred-tank bioreactor. Left panel: OD in logarithmic scale for the full process. Right panel: Determination of specific growth rates (µ) at batch and fed-batch stages by exponential fitting. Each point corresponds to the media of three fermentations. Error bars indicate standard deviations from replicates. **C**. Production parameters in shake flasks and bioreactor. The average and SD of three independent processes are tabulated. Asterisks indicate significant difference at p<0.01 (**) or p<0.0001 (****) in Student’s *t*-tests.

To assess process reproducibility, two additional fermentations were performed in the established conditions until 7 h of fermentation, as at this point we observed the maximum EIT yield. Cultures reached an OD600 of around 7.5 at this fermentation time. The specific growth rate (µ) during the batch stage was 0.57 h^-1^, decreasing to 0.12 h^-1^ during the fed-batch phase up to the time selected for culture collection (7 h), as determined by exponential fitting (Fig. 2B, right panel).

In parallel with each fermentation process, shake flask cultivations were performed in M9Y. Parameters describing EIT production, as EIT concentration, yield and volumetric and specific productivity were determined for both growth conditions (Fig. 2C). EIT concentration and volumetric productivity showed a near 600 % increase when production was carried out in stirred-tank bioreactor compared to shake flasks, while yield and specific productivity presented about 130 % increase. Therefore, a significant improvement was achieved in all parameters (Fig. 2C) indicating a successfully scale-up of EIT production to bioreactor.

### 3. Downstream processing

After 7 h of fermentation, bacteria were collected and subjected to a thermal shock in an appropriate volume of TE buffer. As bacteria can be concentrated around 25-40-fold at this point, there was no need for posterior concentration steps. Upon heat treatment, the periplasmic fraction was recovered in the supernatant.

Samples from both the periplasmic fraction and the pre-treated crude extract were evaluated by Western blot for EIT determination and by Coomassie staining for total protein profile. The same pattern of EIT bands was specifically detected by Western blot after the thermal permeabilization (Fig. 3A, left panel), indicating that EIT was indeed recovered, consistent with previous results (Duarte et al. 2024). However, this procedure does not constitute a purification step, as most protein bands were still detected (Fig. 3A, right panel). Yet, the proportion of EIT in the periplasmic fraction resulted in about twice that of the pre-shock crude extract, as determined by Western blot and Bradford quantitation methods for EIT and total protein content, respectively. Thus, EIT was estimated to represent between 4.0 to 4.8 % of the total proteins present in the supernatant.

**Figure 3.**
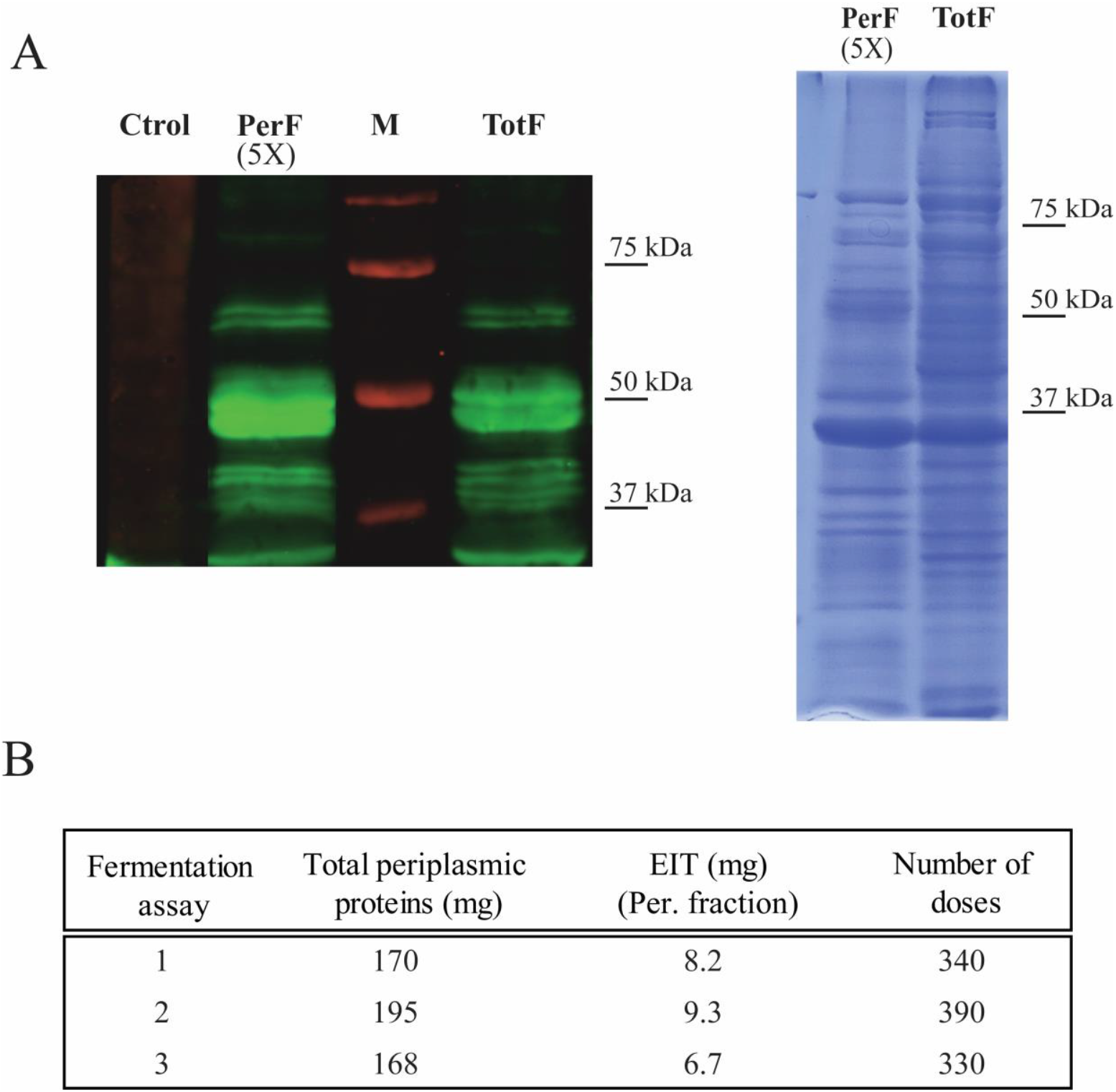
Downstream processing. The periplasmic fraction was recovered after a thermal permeabilization at 7 h of fermentation. **A**. Left, Western blot with anti-EIT antibodies; Right, Coomassie staining. Ctrol refers to *E. coli* carrying the empty vector (periplasmic fraction), PerF and TotF refer to periplasmic and total fractions respectively, with PerF loaded at 5× the concentration of TotF. **B**. Total periplasmic protein content was quantified by Bradford method. Recombinant EIT content in Per fraction was determined by Western blot and ImageJ analysis. BSA or EIT standards were used respectively for calibration. The number of vaccine doses was calculated according to the value of 500 µg of total periplasmic proteins per dose.

In our previous work, we used an EIT-enriched periplasmic fraction derived from *E. coli* cultures in shake flask for a vaccine preparation -upon addition of adjuvant- and tested its immunogenicity in a reduced number of steers (Duarte et al. 2024). Being each dose defined as containing 500 µg of total periplasmic proteins, cultivation in shaken flasks had resulted in the production of 17 doses per L. In the present work, each fermentation yielded an average of 350 doses per L of fermented culture (Fig. 3B), accounting for a considerable improvement of the production process.

### 4. ELISA assays for antigenicity assessment

To test whether the recombinant EIT obtained through fermentation and thermal permeabilization remains functional, we performed ELISA assays using bovine sera previously generated following EIT-immunization (Duarte et al. 2024). By using a purified MBP-EITH7 protein as control antigen, the periplasmic fraction was tested against sera from pre-immune and vaccinated steers corresponding to week 2 post immunization. The periplasmic fraction recovered upon fermentation in bioreactor was specifically recognized by the antibodies of immunized steers, giving OD_450_ values comparable to those obtained with the MBP-EITH7-control, whereas not by the pre-immune serum (Fig. 4). This suggests that the recombinant protein produced by fermentation retains its antigenic properties and is suitable for use in EIT vaccine formulations.

**Figure 4.**
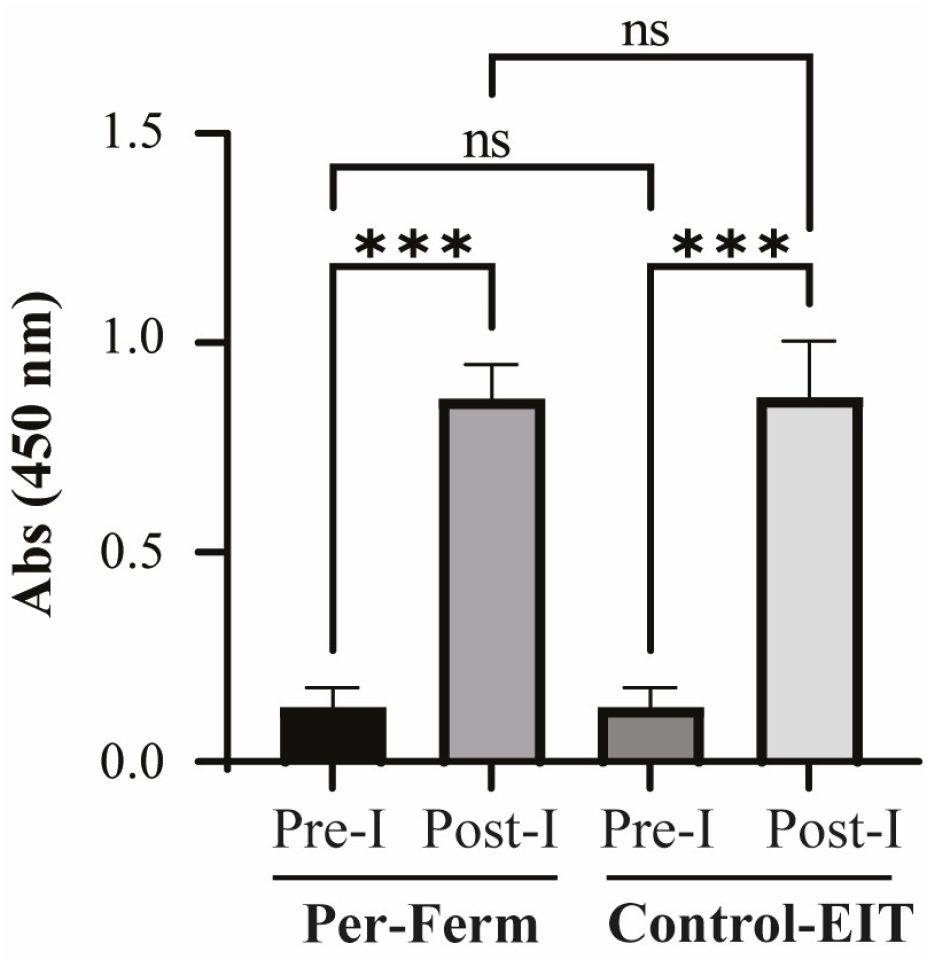
ELISA assays for antigenicity assessment. The recombinant EIT recovered in the periplasmic fraction following fermentation (Per-Ferm) was assayed. Pre-I and Post-I correspond to sera collected before and after steer immunization, respectively. Control-EIT refers to a previously generated purified EITH7 antigen.

## Discussion

Cattle intestinal epithelium is persistently colonized and has been recognized as a major reservoir of EHEC. Bacteria are disseminated into the environment through faecal shedding and food derivatives, and humans acquire infection by oral route. Several strategies have been proposed to reduce the impact of this zoonosis, particularly critical in children below 5-year-old, where EHEC infections can lead to severe sequels and the development of HUS. Preharvest vaccination has demonstrated to be an effective tool for reducing bacterial colonization and environmental spread (Snedeker et al. 2011).

*Escherichia coli* is widely used as host organism for heterologous gene expression. In our recent work, seeking for a simple and cost-effective strategy for a vaccine against EHEC for cattle, we cloned the EIT recombinant antigen fused to a periplasmic signal sequence in the pBBRMCS-2 cloning vector, under a lac-inducible promoter, to be expressed in *E. coli* BL21. As a proof of concept, the periplasmic fraction, formulated with an appropriate adjuvant, was applied subcutaneously and evaluated in both mouse and bovine models, inducing strong antibody response as determined by *in vitro* assays (Duarte et al. 2024). Building on this approach, in the present study we assessed whether recombinant EIT expression could be efficiently scaled up to a laboratory stirred-tank bioreactor, to increase the number of doses obtained in the process, required for more extensive validation studies and potential application.

For each individual product, process and facility, the most relevant process parameters influencing product yield and quality must be determined and set as scale up parameters to be kept as constant as possible (Schmidt 2005). Several factors have been reported to control the bacterial growth and yield of recombinant proteins, as temperature, pH, media composition and additives, inducer concentration, etc. Media composition can be optimized attaining high productivity with short cultivation times and cost reduction. The highest relative quantities of recombinant EIT were detected in minimal medium supplemented with yeast extract (M9Y). In contrast, LB sustained higher biomass but the lowest EIT relative level, while non-supplemented minimal media (M9) resulted in poor bacterial growth. EIT expression was induced by using a lactose-containing solution referred to as GGL5052. Lactose is an economical and non-toxic inducer and, when supplied together with glycerol or glucose -to disfavour its use as a carbon source-it can achieve for efficient recombinant protein expression. Growth temperature was maintained at 37 °C as it has been reported for various recombinant enzymes that *E. coli* BL21 promotes higher expression and yield when growing at this condition (Shahzadi et al. 2021).

Fed-batch fermentations in M9Y produced an EIT concentration of around 30 mg/L and a yield of EIT relative to biomass of 17 mg/g in each process, after 7 h of operation. We found that these parameters decreased during prolonged fermentation. Preliminary results showed faint but detectable EIT bands in the culture supernatants at 6 and 7 h of fermentation (data not shown), suggesting some compromise of the bacterial outer membrane during the process. This unwanted effect could be useful in developing an alternative approach, in which the thermal permeabilization is performed *in situ* in the bioreactor, thus retrieving EIT from both biomass and culture supernatant, increasing EIT recovery. However, an additional concentration step would be required as the culture would not be centrifuged prior to the heat treatment in this strategy.

From a downstream processing perspective, periplasmic expression potentially offers a big advantage. Recombinant protein extraction is facilitated as only the outer membrane needs to be disrupted and, ideally, periplasmic extracts contain only minimal cytoplasmic components (Schimek et al. 2020). However, it has been reported that the common methods described for periplasmic protein recovery and applicable under industrial process conditions, result in enriched fractions but are not truly selective (Schimek et al. 2020). Similarly, by performing a thermal shock in combination with EDTA and Tris, we obtained an extract containing many proteins, in which EIT proportion resulted two-fold enriched compared to the untreated crude extract. Even it is not a purified extract, the periplasmic fraction is intended for direct use in the vaccine preparation. The method is suitable for large scale production and, in addition, drastically reduces the number of viable bacteria upon the thermal shock procedure (Duarte et al. 2024).

In terms of antigenic capability, the immunological assay indicated that EIT produced in stirred-tank bioreactor was comparable to previous antigen preparations, as similar results were obtained when incubated with sera from EIT-immunized cattle.

In summary, we scaled up the production of recombinant EIT antigen by a fed-batch fermentation process in a lab-scale stirred-tank bioreactor, successfully optimizing all production parameters. By using a small volume of around 1 L, the process allowed, in average, the preparation of 350 doses of the vaccine formulation, in a quite simple and cost-effective approach. The fermentation process was performed in triplicate and proved reproducible under the conditions analysed, which is a desirable feature in industrial production processes.

## Acknowledgments

L.A.B., D.G.N., and G.B. are members of the Research Career of CONICET and also serve as faculty members at UNSAM.

## Funding

This work was supported by grant from the CONICET of Argentina (PIBAA 2022-2023 – 28720210100909CO).

## Conflict of interest

None declared.

## Data availability

The data underlying this article are available in Figshare Repository at https://dx.doi.org/10.6084/m9.figshare.30865205

## Notes

### Competing Interest Statement

The authors have declared no competing interest.

